# Establishing an assay for visual desensitization in larval zebrafish for understanding synaptic plasticity

**DOI:** 10.64898/2026.10.28.762667

**Authors:** Oliwia Górska, Limor Freifeld, Kristen E. Severi

## Abstract

Synaptic plasticity, the ability of neurons to modify the strength of their connections, is essential for learning. Desensitization is a type of synaptic plasticity during which a behavioral response to a stimulus is reduced after repeated presentations of a weakened version of the stimulus which does not produce the behavioral response and thus is deemed a subthreshold stimulus. Here we explore whether the larval zebrafish escape response from visual threats, such as an expanding dark spot, can act as a potential model to study desensitization. We developed a novel experimental setup to investigate visual desensitization, exposing larvae to looming visual threats of varying contrast, and used high-speed cameras to verify rapid escape movements. First, we established stimuli which reliably acted as subthreshold and suprathreshold stimuli. Next, we quantified the response rate before and after desensitization and compared those data to control larvae who were not desensitized. We observed desensitized larvae reduced their response rate to threatening looming stimuli. By providing an opportunity to explore the plasticity mechanisms governing sensory-motor transformations and behavioral adaptation, this research contributes to our understanding of fundamental principles of learning and neural plasticity across vertebrates.

## Introduction

All animals must correctly interpret sensory inputs to choose behavioral responses that maximize their survival chances. Neural circuits mediating sensory-motor transformations implement these behavioral choices and must be adaptive to allow modulating behavior by experience. Sensory experience can vary between multiple modalities in converging or competing paths and yet must be used to inform a behavioral choice. The circuit mediating escape responses to auditory, visual or tactile stimuli, in teleosts, including zebrafish, has long been a model of decision making and synaptic neurobiology (**Figure 1A**; Zottoli and Faber, 2000; Korn and Faber, 2005; Medan and Preuss, 2014; McArthur et al., 2020). The fast, C-start escape response is mediated by the firing of a single action potential in one of the two Mauthner cells, a pair of large “command”-like neurons residing in the hindbrain (Hale et al., 2016). The Mauthner cells excite primary motor neurons and interneurons in the spinal cord (Berg et al., 2018), bending the fish body into a C-shape and turning it away from the stimulus source (Kimmel et al., 1974). Which of the two Mauthner cells is activated and the timing of this activation are important for survival, and are gated by a well investigated network of interneurons (Koyama et al., 2011; Koyama et al., 2016; **Figure 1B**). Nevertheless, escape responses with different timing and kinematics can still occur if both Mauthner cells are activated bilaterally or not at all (O’Malley et al., 1996; Liu and Fetcho, 1999; Bhattacharyya et al., 2017; Liu and Hale, 2017; Marquart et al., 2019). Activity imaging in the Mauthner cells and their segmental homologous in response to threatening stimuli (Gahtan et al., 2002; Bhattacharyya et al., 2017), as well as ablation studies (Liu and Fetcho, 1999; Dunn et al., 2016) indicate that the motor circuit gives rise to flexible escape responses triggered by multiple sensory modalities. Escape responses can be induced by auditory and visual stimuli: startling acoustic stimuli activate auditory fibers of the VIIIth cranial nerve forming excitatory mixed electro-chemical synapses with the lateral dendrites of ipsilateral Mauthner cells, eliciting escapes within a few milliseconds of stimulation (Eaton et al., 2001; Yao et al., 2014) while startling visual stimuli require a longer pathway, and arrive via the optic tectum to Mauthner cell ventral dendrites (Zottoli et al., 1987; Sato et al., 2007; Helmbrecht et al., 2018; **Figure 1**).

**Figure 1.**
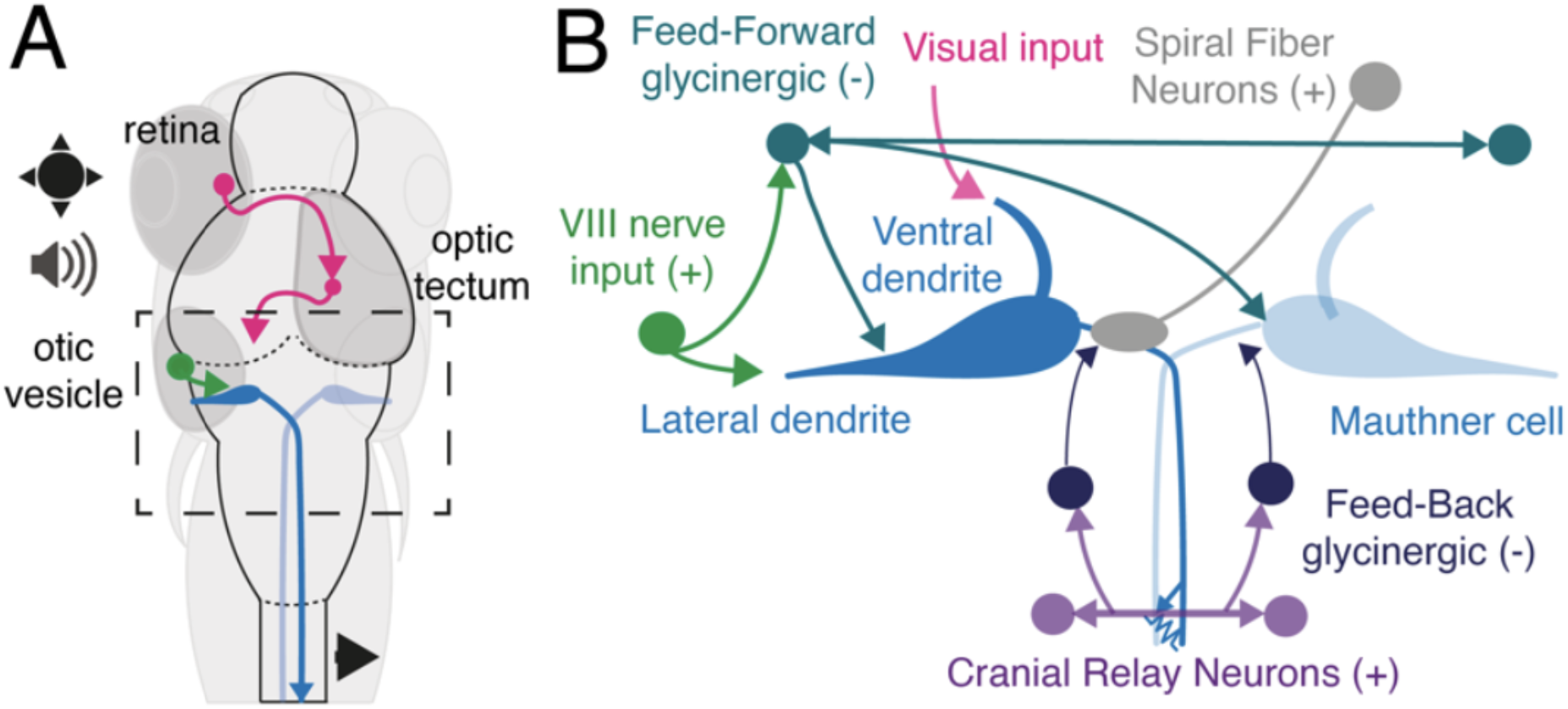
Components of the escape circuit in larval zebrafish. A. Acoustic (green) information arrives at the Mauthner cells via the VIII cranial nerve while visual signals (pink) arrives from the retina through the optic tectum. **B**. Circuit components initiating and gating Mauthner cell activation (box in A). Auditory information targets the Mauthner cell lateral dendrite while visual information targets the ventral dendrite. Feed-Forward (FF) inhibitory glycinergic neurons target both Mauthner cells and their contralateral population, while excitatory Spiral Fiber Neurons enhance activity at the Mauthner cell axon cap. Cranial Relay Neurons are activated by the passing Mauthner cell axon and excite bilateral populations of Feed-Back (FB) glycinergic interneurons targeting the Mauthner cells.

The way the decision to escape is implemented and modulated by experience is better understood for auditory inputs. Repetitive presentation of subthreshold auditory stimuli causes desensitization, a decrease in the fast C-start escape response probability (Oda et al., 1998; Takahashi et al., 2017), via strengthening of glycinergic synapses on the Mauthner cells (Ogino et al., 2019). However, it is unknown whether visual desensitization occurs and if so, if it occurs via the same mechanism. To explore this question, we tested whether plasticity in the escape response could occur through the visual circuit using a behavioral response as a readout. We hypothesized that when exposed to suprathreshold stimuli following repeated subthreshold stimuli in an effective desensitization paradigm, larval zebrafish would reduce their response rate to suprathreshold looming stimuli compared to control animals and initiate fewer escape responses.

## Methods

### Animals

Larval zebrafish used in these experiments were 6 days post-fertilization (dpf) AB Wild-Type (origin: ZIRC stock center, Eugene, Oregon). They were reared in an incubator at 28.5°C with a 14L:10D light cycle. Larvae were generated from an adult colony maintained at the New Jersey Institute of Technology in the Severi lab under Rutgers University-Newark IACUC oversight, PROTO201800041.

### Behavioral setup

High-speed videos were collected on a custom-built setup (Zaki et al., 2021). A high-speed camera (Mikrotron GmbH, Germany) fitted with a 35 mm F1.4 lens (Fujinon) was used to acquire images.

Larvae were additionally illuminated with 850 nm IR LEDs which are not within their visible spectra (Waveform Lighting) under an acrylic platform covered with light diffusers (Pro Gel, B&H Photo) within a custom-built enclosure (MiniTec Framing Systems, LLC) which was left closed throughout the experiments. Visual stimuli were rendered with OpenGL and expanded from the center of the dish linearly over the course of 300 ms to a final diameter of 5 cm and then remained fully expanded for an additional 300 ms (**Figure 2A**). Visual stimulation was presented in open-loop via a small projector (3M) and reflected off a cold mirror (Knight Optical) located under the platform and above the IR LEDs at a roughly 45 degree angle. Videos were acquired at 250 Hz with 1,423 μs shutter speed at 648 × 648 pixel resolution and saved as AVI format using custom-written LabView software (National Instruments). Both ‘pre’ and ‘post’ videos were 75,000 frames and 300 s in duration. ‘Pre’ and ‘post’ trials used the same individual larvae, and each larva was tested one at a time. The videos were captured 5 minutes after larvae were placed in the behavior enclosure in order to allow time for acclimation following handling. Animals were recorded at a temperature of 25-28°C in round petri dishes with 5.0 cm diameter.

**Figure 2.**
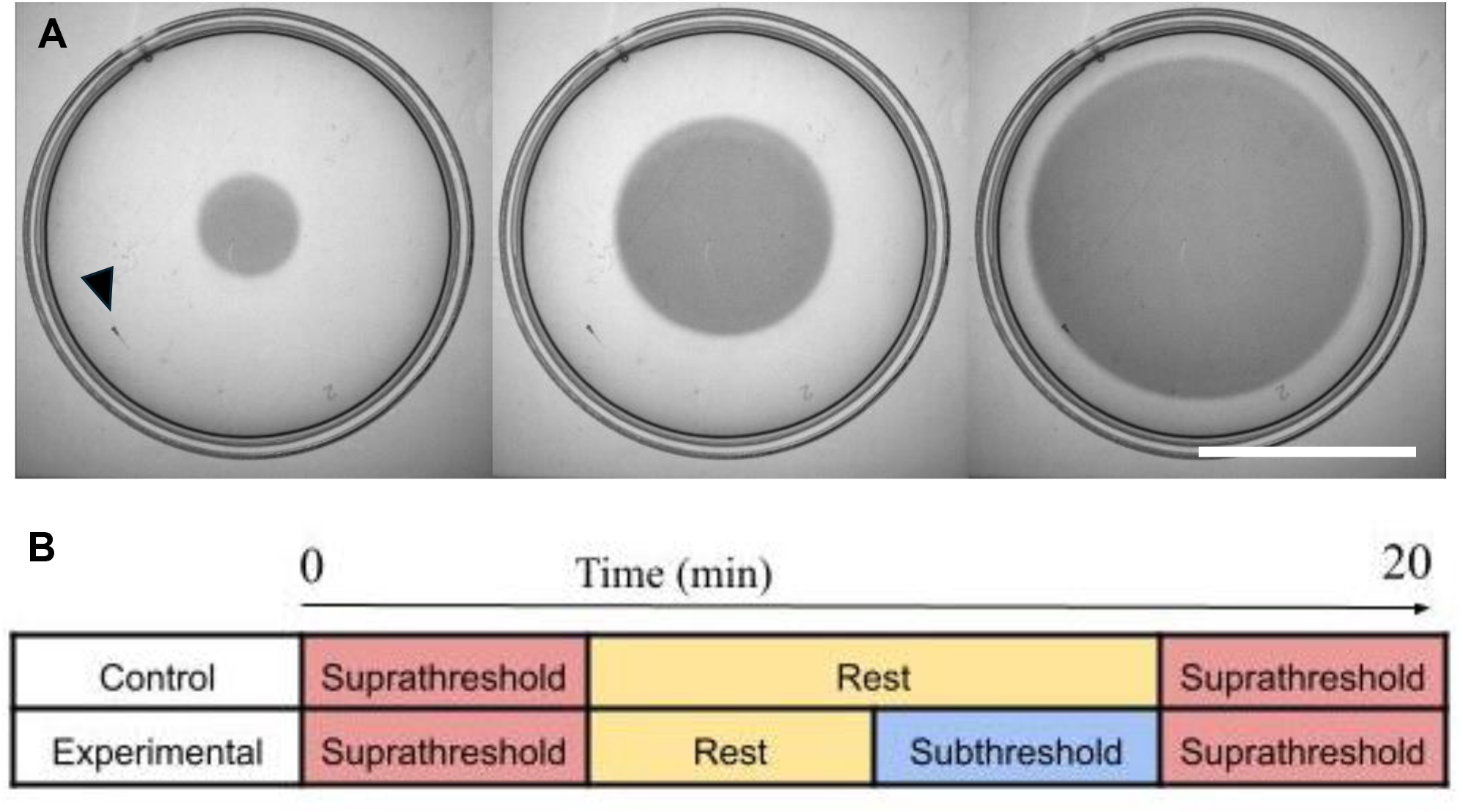
Experimental protocol. A. Raw images with 100 ms between frames during looming stimulation. Looms were positioned to appear from the center of the dish each time. Larval position is indicated by arrowhead. Scale bar is 2.5 cm. **B**. Protocol schematic. One block represents 5 minutes. During the suprathreshold stimulation period there is a stimulus shown every 30 seconds (10 looms total). During subthreshold stimulation period there is a stimulus shown every 5 seconds (60 looms total). During the rest period there is a white screen. Videos are acquired during the suprathreshold presentation periods.

### Stimulation protocol

The experiments were divided into two groups: control and experimental treatments. Both groups had the same camera acquisition settings and duration, 20 minutes in total, but had different stimulation procedures (**Figure 2B**). Control larvae were exposed to 5 minutes of suprathreshold stimulation which consisted of an expanding spot at full contrast (black on a white background) appearing once every 30 seconds, as the pre-exposure period. The control larvae were then left undisturbed with the white background for 10 minutes. The suprathreshold stimulation was then presented for 5 minutes in an identical procedure in the post-exposure period. Experimental larvae were exposed to an identical period of suprathreshold stimulation, followed by only 5 minutes of rest. They were then exposed to 5 minutes of subthreshold stimulation, which included a gray spot (contrast reduced to 60% of original) on a white background. We determined that this was the minimum contrast reduction that reliably never induced visual escapes. The size and expansion rates of the two looming stimuli were identical, and different only in contrast. During the subthreshold stimulation, low contrast looms were shown in rapid succession, every 5 seconds, for 5 minutes. This was followed immediately by a suprathreshold stimulation block that constituted the post-exposure period. Video was recorded only during the suprathrehold periods (both ‘pre’ and ‘post’) to determine the response rates for each period. Response rates were determined from reviewing video manually and confirming canonical kinematic signatures of escape such as a C-start followed by a burst swim (Budick and O’Malley, 2000) as present or absent during each loom presentation period.

## Results

A black spot expanding on a white background under closed-loop conditions evokes escape responses (Dunn et al., 2016; Heap et al., 2018). Using this as a guide, we have developed a desensitization protocol for open-loop visual looms modeled on established auditory desensitization protocols (Oda et al., 1998; Takahashi et al., 2017; Ogino et al., 2019). We utilized a low contrast grey spot on a white field looming stimulus which was otherwise identical to our full contrast looming stimulus. This low contrast version reliably produced a lower response rate (1/50 or 2% response rate, n = 10 larvae) compared with full contrast stimuli (19/60 or 31.7%, n = 12 larvae), and so we deemed this an appropriate subthreshold stimulus (Temizer et al., 2015; Yao ett al., 2016; **Figure 2**). We viewed each recorded video and confirmed the kinematic signatures of an escape response (**Figure 3A**) to determine whether the movements constituted escape behavior or simply routine turns, slow swimming, or other navigational maneuvers which would not be consistent with escape (Budick and O’Malley, 2000; Burgess and Granato, 2007; Marques et al., 2018).

**Figure 3.**
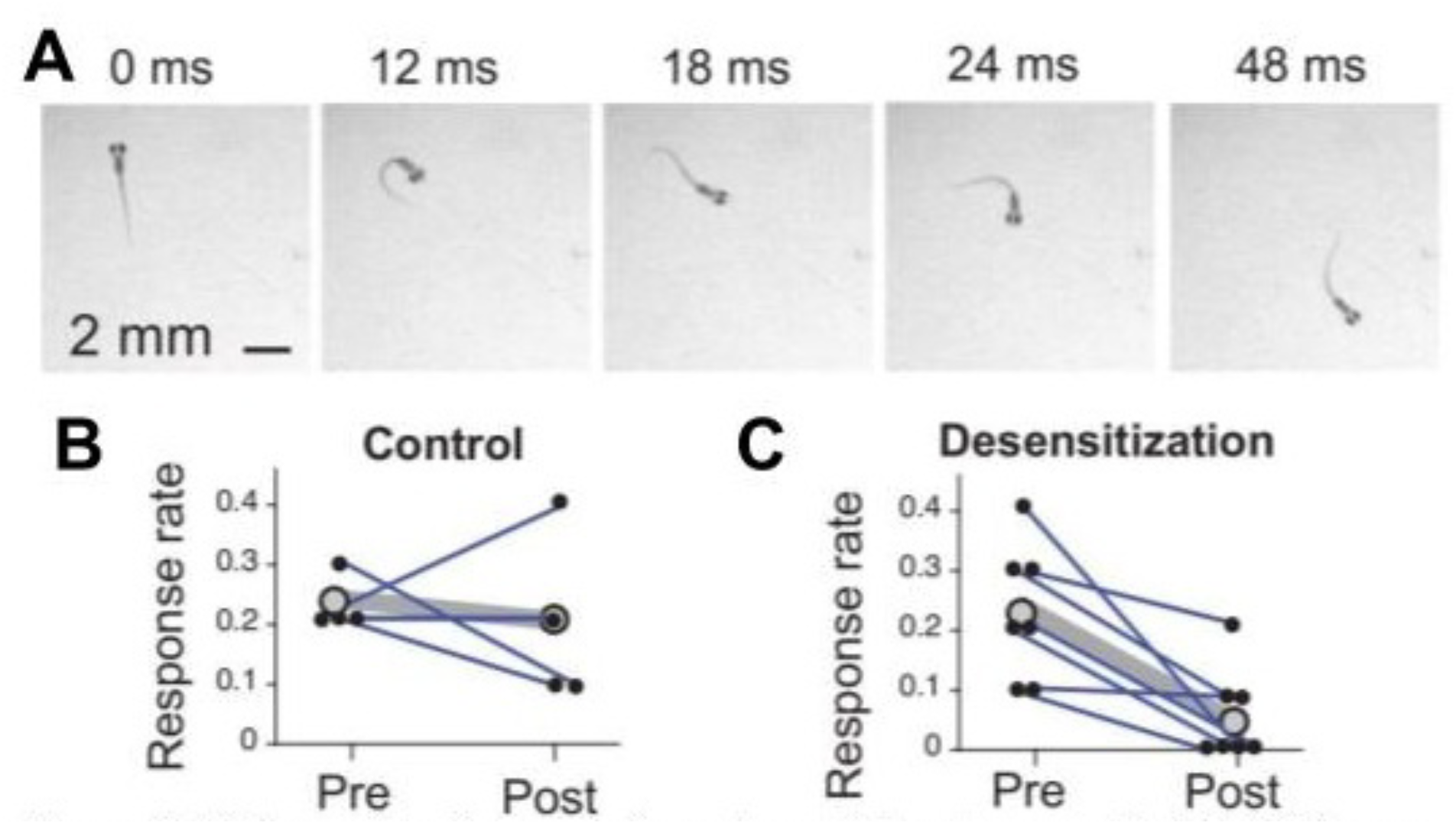
Visual desensitization induced in larval zebrafish by presenting low contrast looms. **A**. Raw video frames during a loom-induced escape trial demonstrating canonical escape behavior. **B**. Escape response rates to full contrast looms before and after a rest period (control). **C**. Escape response rates to full contrast looms following a visual desensitization period using reduced contrast looms. **B-C**. Blue lines match ‘pre’ and ‘post’ stimulation for the same individual (black). Grey dots are the average per condition with the grey line indicating the matched averages across ‘pre’ and ‘post’ conditions. N= 4 control larvae and 7 desensitized larvae. Each larva was given 10 pre and 10 post presentations.

Having established the efficacy of our visual stimuli, we proceeded to assess whether repeated exposure to subthreshold stimuli would decrease the response rate to full contrast suprathreshold stimuli, a hallmark of desensitization (Ogino et al., 2019). Indeed, desensitization to looming stimuli did occur, with the exposure to subthreshold stimuli immediately preceding suprathreshold presentations decreasing the response rate from an average of 23% to an average of 6%, whereas control larvae who were not exposed to the desensitization protocol maintained an average response rate of 23% (pre) and 20% (post) (**Figure 2B-C**).

## Discussion

Mauthner cells take in sensory information from either side of the body and elicit a strong contralateral contraction of body muscles allowing for an evasive turn (Bhattacharyya et al., 2017). Visual stimuli activate the Mauthner cells after processing in the optic tectum and elicit directional responses away from the stimulus (Dunn et al., 2016). The looming stimulus utilized here was an expanding black spot on a white field, presented from below, and increased in size to simulate a rapidly approaching threatening object (Bhattacharyya et al., 2017). The Bhattacharyya study focused on the habituation, a decrease in response magnitude to repeated presentations of an identical stimulus, a well-known and important form of synaptic plasticity in many organisms including zebrafish (Burgess and Granato, 2007; Wolman and Granato, 2012; Köcher and Straumann, 2025). Here we focused on desensitization, a similar form of synaptic plasticity during which repeated presentation of a weaker, subthreshold version of the stimulus reduces the response to a strong, suprathreshold version of the stimulus. Desensitization at the Mauthner cell has been shown using acoustic stimulation (Ogino et al., 2019), and we based our protocol in part on this study, but using the visual modality. Here we demonstrate this form of plasticity in larval zebrafish for the first time.

We observed some limitations of the open-loop presentation paradigm, which resulted in a relatively low response rate in both conditions as the larva was not always positioned identically in the dish or at a fixed distance from the spot centroid at the time that the looming spot appeared. While the larvae did at times perform escape responses on the periphery of the dish, they often did not respond when in direct contact with the wall. A closed-loop setup would likely be more effective (as in Dunn et al., 2016; Marques et al., 2018).

The objective of this study was to open a door to our understanding of desensitization, which can be investigated uniquely effectively in the larval zebrafish model. The establishment of this paradigm will provide an opportunity to compare sensory-specific or interacting neural circuit modulations, both at the level of synapses as well as the level of recruitment of the glycinergic inhibitory populations. Auditory desensitization is understood to be accomplished mechanistically via synaptic changes at glycinergic synapses onto the Mauthner cell (Ogino et al., 2019). Glycinergic synapses found on larval zebrafish Mauthner cells are similar to mammalian synapses in their structure and their function, and as such, mechanisms that impact their plasticity and sensory-motor transformation to alter behavior may be suggestive of mechanisms common to all vertebrates (Betz and Becker, 1988; Legendre et al., 1995; Hirata et al., 2005; Ogino et al., 2019).

## Acknowledgements

We thank Dr. Alfred Amendolara for his excellent contribution toward rendering the loom visual stimuli and the lab of Dr. Michael Orger for sharing OpenGL code for visual stimulation. We thank the Zebrafish International Resource Center for providing AB WT fish, and the Severi Lab for zebrafish husbandry support. O.G. was supported by an NJIT Provost Undergraduate Research Innovation Fellowship. L.F. was supported by the Chief Scientist Office (CSO) of the Israeli Ministry of Health (MOH) as part of an ERA-NET co-fund scheme ERA PerMed, and by the Zuckerman STEM Leadership Program. K.E.S. was supported by NJIT Startup funds.

